# No Relationship Between Perceived Health Anomalies and Perceived Experimental Success in Retired Breeder Male Hartley Albino Guinea Pigs

**DOI:** 10.1101/2020.03.06.979336

**Authors:** Chandra B. Bain, Julie M. Settlage, Grace A. Blair, Steven Poelzing

**Affiliations:** Department of Biomedical Engineering and Mechanics, Virginia Polytechnic Institute and State University, Blacksburg, VA; Fralin Biomedical Research Institute at Virginia Tech Carilion, and Center for Heart and Regenerative Medicine, Roanoke, VA; Translational Biology, Medicine and Health graduate program, Virginia Polytechnic Institute and State University, Blacksburg, VA; Office of the University Veterinarian, Virginia Polytechnic Institute and State University, Blacksburg, VA

## Abstract

Guinea pigs used in our laboratory for cardiac research sometimes exhibit physical abnormalities. These issues may abate or intensify during the time they are housed in our facility. After using a guinea pig for research, experimentalists note the apparent health of an animal based on visible features and/or abnormal electrophysiology of the heart. There was an existing anecdotal observation that the health of the Guinea Pigs, and subsequently the experimental success rate, had a seasonal variation; therefore we sought to determine if there is a time of year in which our guinea pigs are more likely to be perceived as unhealthy, and whether any determined monthly pattern correlates with an experimentalist’s ability to complete an experimental protocol. An electronic log was created to record the perceived health of the animal and the ability to complete the experiment successfully. Irregular symptoms included, but were not limited to, severe weight or hair loss and irregularities with the heart found post thoracotomy or during baseline electrophysiological recordings of whole-heart preparations. Animals that did not exhibit significant weight or hair loss, or other ailments were considered “healthy”. Overall, our results indicate that there are no monthly variations in perceived Hartley Albino guinea pig health or correlations with experimental completion rates, suggesting mild hair or weight loss that is common when shipping animals may not significantly affect the ability to conduct *ex vivo* whole-heart electrophysiological studies.

## INTRODUCTION

Guinea pigs are useful as models in cardiac research over rats or mice^1^ because their electrophysiological profile more closely resembles those of larger animals.^2^ Our group at the Fralin Biomedical Research Institute at Virginia Tech Carilion (FBRI) studies the electrophysiology of the isolated guinea pig heart in order to elucidate mechanisms that can conceal or unmask cardiac conduction related disease. Animals used for our research studies are shipped from the vendor weekly and housed on site for typically one week or less before they are studied. During this short time in the vivarium, the animals receive daily health checks and weight monitoring, rarely requiring more extensive monitoring. Still, experimentalists may pass along to junior colleagues their perception that the likelihood of completing an experimental protocol is seasonally dependent, and that this might be evident by visible aberrations with animals. Therefore, we wanted to prospectively determine whether a correlation exists between animal mortality, perceived animal health, month and likelihood of completing a Langendorff-perfused guinea pig whole-heart experiment using male retired breeder Hartley albino guinea pigs. We tested the null hypothesis that there was not a significant relationship between time of year and occurrence of perceived physical irregularities. We further sought to determine whether perceived animal health (hereafter referred to as healthy or unhealthy) affects our perceived experimental success rate (hereafter referred to as success).

## MATERIALS AND METHODS

This study was carried out in accordance with the principles of the Basel Declaration and recommendations of the *Guide for the Care and Use of Laboratory Animals,* National Research Council.^3^ The protocol was approved by the Institutional Animal Care and Use Committee (IACUC) at Virginia Polytechnic Institute and State University.

### Animals

In addition to demonstrating an electrophysiological profile similar to humans, the Hartley guinea pig strain is the most commonly used guinea pig for research.^4^ To include other strains besides Hartley albino could introduce more genetic variability. Similarly, using only males removes any variability between the sexes. The risk of cardiovascular disease increases with age^5^ while animal purchase costs decrease with age. By using retired breeder guinea pigs we incorporate an aged-animal model while reducing the number of guinea pigs bred by the vendor and decreasing our animal purchase costs. Male retired breeder Hartley Albino guinea pigs (N=1338, 697-1543g, 12-15 months old) were purchased from Hilltop Laboratory Animals, Inc., Scottdale, PA. Animals were shipped in a 24 x 17 x 7 inch animal-safe carton. Each carton contained at most two guinea pigs with a divider in between them to maintain physical separation. Based on health surveillance performed by Hilltop, all guinea pigs were free from Lymphocytic Choriomeningitis Virus, Pneumonia Virus of Mice, Reovirus 1, 2, 3, Sandai Virus, Guinea Pig Adenovirus, Bordetella, helminthes, pathogenic protozoa, ameba and ectoparasites. Primary bacterial pathogens were not detected by aerobic culture techniques. Guinea pigs were held in an animal biosafety level 1 project designated housing-room until needed for research.^6^ Guinea pigs were housed one to a cage and under the ideal environmental conditions of 20-26°C and 30-70% humidity.^7, 8^ Animals were exposed to twelve-hour light and dark cycles. Fresh air was pumped into the containment room. Hay, a red-colored shelter and a toy were placed in the cages for diet and enrichment, respectively. Animals had access to Vitamin C supplemented (0.5 mg/gm) standard guinea pig chow ad libitum.^9^ Reverse Osmosis water was offered in a standard cage water bottle that differs from the apparatus they used in their origin facility. If boarding lasted longer than one-week post-delivery, cages were changed for clean ones.

### Langendorff-Perfused Heart Preparation

Oscar Langendorff established the Langendorff technique in 1897 as an *ex vivo* model to study the mammalian heart. This whole-heart model provides a broad platform for our research areas of interest.^10^ At the beginning of each experiment Isoflurane (3-5% at 3-4 L/min oxygen flow) or sodium pentobarbital (Nembutal, 30mg/kg IP) was administered until the animal was insensate following approved protocols. Hearts were then removed by thoracotomy and retrogradely perfused in a Langendorff preparation with oxygenated variable HEPES-buffered solution and the atria removed to reduce competitive stimulation.^11^ Heart irregularities witnessed during the initial baseline readings were noted. However, aberrations including but not limited to a wide QRS complex, infarctions or asystole found during any administered perfusate other than our Lab Standard Control were excluded from perceived health data analysis. Similarly, any irregularities found once any experimental interventions were introduced (other than our routine administration of the membrane potential fluorophore, Di-4-ANEPPS) were excluded from the perceived health data analysis.

### Statistical Analysis

Multiple IACUC approved active laboratory personnel collected data over a period of fifty-two months on the research protocol. Experimentalists noted their overall perception of the animal’s health and their success in completing their experimental protocol in an electronic database after each experiment. Health was a binary variable, where a mark of “1” indicated the experimentalist perceived the animal and/or heart was unhealthy and “0” indicated a healthy animal with a freshly isolated heart free from irregularities. Experimental success was also a binary variable where a “0” denoted a successful experiment, in which the protocol was completed successfully, and a “1” denoted an unsuccessful experiment where the protocol was terminated prematurely. A protocol may have been terminated prematurely due to a variety of reasons including but not limited to problems with the heart or operator error. Researchers were then asked to fill in the comments section with details about the perceived abnormality and completion of the protocol where appropriate. The authors reviewed all comments for clarity and equalized results as needed. Statistical analyses were performed using one-way Analysis of Variance (ANOVA) tests or linear regression analyses of unpaired data. A p-value less than or equal to 0.05 was considered statistically significant to reject the null hypothesis. All values are reported as mean ± standard error of the mean unless otherwise noted. Graphpad Prism or Microsoft Excel software were used to conduct statistical analyses.

## RESULTS

### Experimental Numbers and Perceived Health

Figure 1A depicts our average monthly counts for numbers of animals used in research. Importantly, experimental load over the past four and a half years did not vary significantly during the course of the calendar year. Experimentalists were asked to judge whether an animal or heart was unhealthy after conducting an experiment and noted this in laboratory notebooks and the electronic database. Irregular health by visual inspection of the animal prior to surgery included hair or weight loss affecting 10% or more of the body, lethargy and death in housing. Once the heart was extracted, an animal was deemed unhealthy if a visual inspection found abnormalities including but not limited to fibrotic scarring, a calcified aorta or an enlarged or discolored heart. Finally, since studies in our laboratory focus on abnormal electrophysiology and arrhythmias, isolated hearts were considered unhealthy if they exhibited a wide QRS complex measured from a volume-conducted ECG >35 msec during intrinsic rhythm or the if heart went into spontaneous VF or VT (Ventricular Fibrillation/Tachycardia) during the initial equilibration period prior to experimental data collection. There were a few incidences of infarction, asystole, ischemia and fatty liver recorded. Rarely, issues such as an unidentifiable mass, calcification of the cartilage associated with the sternum, skin problems or fat on the left ventricle or coronary artery were noted. Of 1338 animals inspected in four and a half years, twelve died from natural causes (less than 1%) and 793 exhibited one or more of the above irregular symptoms and were adjudged to be unhealthy (59%). Significant hair and weight loss events, when combined, were responsible for 55% of health irregularities reported.

**FIGURE 1.**
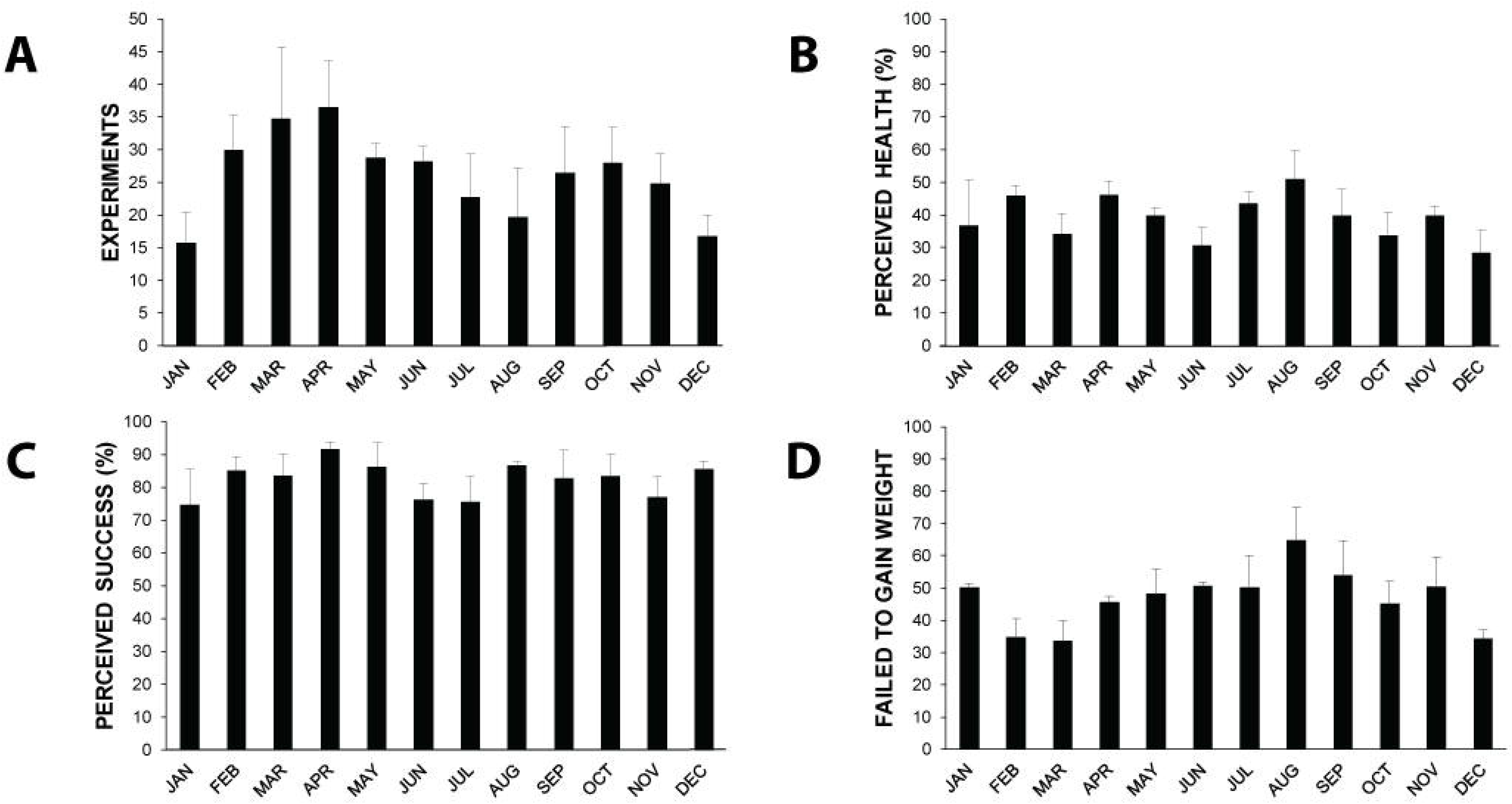

Additionally, a one-way ANOVA of perceived health monthly averages alone throughout the calendar year (Figure 1B) does not reveal significant differences. This suggests there is no associated risk by month for animal health throughout the calendar year. It is important to note that the ANOVA may not have revealed a significant difference because there were only four data points per month corresponding to each year of this study (with the exception that June and October-December data included five years).

### Experimental Success Rate

Before relating the perception of animal health to experimental success, it is important to assess our own track record for experimental success, independent of experimental health bias. The percentage of perceived successful Langendorff whole-heart experiments throughout the calendar year can be found in Figure 1C. Overall, the laboratory averages an 82% experimental success rate, demonstrating a relatively high level of proficiency that may be independent of animal health. A one-way ANOVA revealed no significant difference in our monthly averages of successful experiments.

### Weight Loss

It is common for aged guinea pigs to lose weight during shipping due to stress or neophobia.^12^ We found that only six out of 1,271 animals (0.5%) gained weight during shipping. Animals were weighed before leaving Hilltop, within 24 hours of arriving at the FBRI, and once daily for the duration of their housing in our vivarium. Once a guinea pig lost greater than or equal to five percent of their arrival weight (weight at the time of intake at FBRI), the standard operating procedure at FBRI required a warmed 0.9% Sodium Chloride (saline) solution to be injected subcutaneously once daily as needed. Volume of saline injected depended on animal weight at the time of arrival. Roughly 24 hours after injection, the vivarium staff and university veterinarian determined whether further saline injections were necessary based on current weight and whether or not the guinea pig was eating and drinking on their own. Ideally, all animals would have gained weight after a short acclimation period. However, this was not always the case and only 56% of the animals who lost a gram or more of weight during transportation regained weight after an acclimation period of at least 24 hours and up to fifteen days. Figure 1D suggests that animals were less likely to gain weight during August however a one-way ANOVA reveals no significant differences amongst monthly averages.

### Parametric Relationships

To directly ascertain if months with a larger experimental load were associated with a perceived health bias and as a result transform small differences into statistically significant differences, a bivariate analysis of these two variables was performed, and the data is presented in Figure 2A.^13^ No positive correlation was found between the average number of monthly experiments and animal health. This suggests there is not a significant animal health bias when more animals are used in a given month for research. Likewise, a bivariate analysis of health and experimental success rates (Figure 2B) fails to reveal a strong linear relationship. These data suggest that there is no correlation between our perceived animal health issues and our experimental success rate.

**FIGURE 2.**
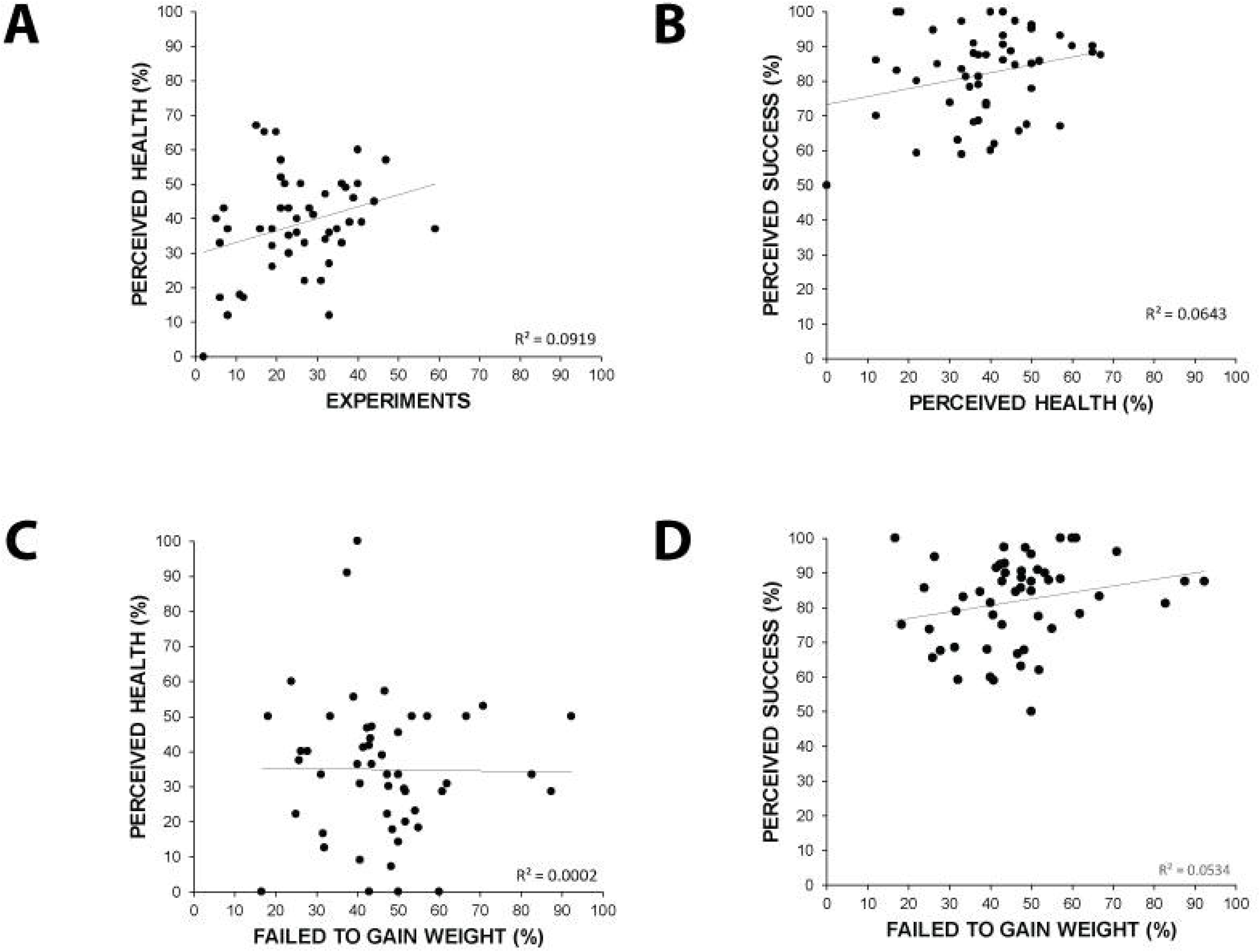

We wanted to resolve if failure to regain weight post shipping was a precursor to significant weight loss (defined here as ≥10% of shipped body weight) or any of the other irregular health events mentioned previously. Here health data only includes those animals who did not gain weight after some period of acclimation. A linear regression analysis (Figure 2C) fails to support a reproducible relationship. The data lead us to reason that an animal who failed to regain weight after shipping is not more likely to experience significant weight loss or other symptoms including but not limited to hair loss, lethargy or natural death.

To see if a non-biased measure like weight loss is a better predictor of experimental success, we performed a bivariate analysis of the two in Figure 2D. An R^2^ value of 0.05 suggests there is no linear relationship between a failure of an animal to regain weight post shipping and experimental success.

## DISCUSSION

These data were analyzed to investigate the relationship between aged guinea pigs perceived as being unhealthy and *ex vivo* electrophysiology experimental success. In support of our hypothesis, we conclude that there is no statistically significant difference in the health of Hartley Albino guinea pigs during any particular month of the calendar year. Fortunately, there is no strong correlation between animal health and success of our electrophysiology research. By reviewing the additional data collected, we are able to show our proficiency with the Langendorff technique as well as our monthly number of guinea pigs used for research. Importantly, we are not introducing a strong bias during our most productive months. In addition, we were able to show that in our time frame, weight loss post shipping was not a significant precursor to irregular health symptoms. From a cardiac electrophysiology standpoint, we feel the success data should reassure other investigators that any of our listed health aberrations seen in male retired breeder Hartley Albino guinea pigs are statistically negligible for electrophysiology research.

These results are important for future studies. Investigators and animal facility managers who have seen similar irregular but not inhumane aberrations in their guinea pig colony may find our list of observed health anomalies useful or reassuring. Investigators requesting an animal protocol may want to account for a one percent loss in total animal numbers due to our reported mortality rate for male retired breeder Hartley Albino guinea pigs. Investigators performing studies on the skin, hair or weight of guinea pigs can use these data to contemplate if male retired breeder Hartley Albino guinea pigs are the best model for their research and readjust their protocols before any animals are sacrificed. While no environment is entirely stress-free, the IACUC and animal vendors can use our listed aberrations to improve current shipping and living conditions of male retired breeder Hartley Albino guinea pigs. Alleviating some travel-induced stress could reduce the number of animals exhibiting significant weight or hair loss, which may prevent exclusion from a study based on experimentalist’s visual inspection and ‘a priori’ rejection of the animal.

### Limitations

One limitation of this study is user bias. Visual health assessments were conducted by 12 different investigators. As these assessments were largely subjective, it is possible that one investigator may conclude hair loss, for example, was worth recording while their colleague may not agree. Different researchers may conclude that an experiment was a failure or a success at varying non-specified time points within their experimental protocol. Another limitation of this study is that it is hard to determine how much of the weight loss we see post shipping is due to inherent differences in the Hilltop versus FBRI scales and how much is actually due to stress. Weight differences between Hilltop and FBRI scales range greatly from 0.78% to 24.81% difference with an overall average of 8% difference since the beginning of this study.

It is worth noting that over the past couple of years we also tried various interventions to relieve stress and improve acclimatization post travel. These included providing the animals with drinking water obtained from Hilltop Labs, feeding them apple slices to assist with water intake or placing some used female bedding in the cage with the male upon arrival at FBRI. None of the changes were deemed to be beneficial by the staff observing the animals, however the specific time period of each intervention wasn’t digitally recorded and cannot be analyzed statistically.

In addition, human error during data entry cannot be ruled out. Our laboratory has two optical mapping rooms, which are both subject to equipment failures that may or may not affect the physiology of the heart. Also, another lab may use a different control perfusate and voltage-sensitive dye and therefore may find fewer or more abnormalities with the heart during baseline readings. One more limitation is that for six out of the fifty-two months data was collected, animals were anesthetized with sodium pentobarbital instead of Isoflurane. In this paper, we do not break down our data by anesthesia treatments. While it is unlikely, any variability introduced by type or dosage of anesthesia must be acknowledged as a limitation to this study.

## Acknowledgments

Many thanks to current and previous lab members for diligently recording details of their experiments for analysis. The authors would like to acknowledge Dixon Smiley and Jenny Raines as former FBRI Vivarium Managers and current vivarium staff for their support with this manuscript. The authors would also like to thank Dr. Gregory Hoeker for his expert input and technical writing assistance with this manuscript.

