## Supplementary material for "No Relationship Between Perceived Health Anomalies and Perceived Experimental Success in Retired Breeder Male Hartley Albino Guinea Pigs": Figure Legends

Figure 1. (A) Number of Experiments by Month. One-way ANOVA reveals no significant differences in monthly average number of experiments (p= 0.3042). (B) Perceived Health by Month. One-way ANOVA reveals no significant difference in reported health symptoms relative to calendar month (p = 0.4269). (C) Perceived Success by Month. One-way ANOVA reveals no significant difference in success rate (p= 0.7050). (D) Failed to Gain Weight by Month. One-way ANOVA reveals no significant difference in a lack of weight gain by month (p= 0.1289). Data represents monthly averages over approximately four and a half years for all graphs depicted above.

Figure 2. Bivariate analyses. (A) Health and Number of Experiments. Linear regression analysis reveals there is no correlation between animal health and the number of experiments conducted each month as evidenced by an R2 of 0.09. (B) Success and Health. Analysis of health affecting experimental success. As evidenced by an R2 value of 0.06, this data does not fit a linear regression model. (C) Health and Weight Loss. A failure to gain weight is the predictor variable and health is the response variable. A linear regression reveals no direct correlation between a failure to gain weight and animal health as evidenced by an R2 value <0.01 (D) Success and Weight Loss. A linear regression analysis reveals there is no direct linear relationship between a failure to gain weight and the success of an experiment as evidenced by a R2 value of 0.05. Data points are monthly averages over 52 months.
